# Interplay between Hyperglycemia and Chikungunya Virus Infection: Pathophysiological Insights from Murine Model

**DOI:** 10.1101/2025.08.01.668212

**Authors:** Pedro Henrique Carneiro, Tháyna Sisnande, Bia Francis Rajsfus, Matheus Silva de Souza, Mariana Juliani do Amaral, Lucas Mendes-Monteiro, Leandro Miranda-Alves, Kissila Rabello, Caio Gonçalves Azevedo, Jorge José de Carvalho, Diego Allonso, Ronaldo Mohana-Borges

**Affiliations:** Division of Infectious Diseases and Vaccinology, School of Public Health, University of California, Berkeley, Berkeley, CA, USA; Laboratório de Biotecnologia e Bioengenharia Estrutural, Instituto de Biofísica Carlos Chagas Filho, Universidade Federal do Rio de Janeiro, Rio de Janeiro, Brazil; Departamento de Biotecnologia Farmacêutica, Faculdade de Farmácia, Universidade Federal do Rio de Janeiro, Rio de Janeiro, Brazil; Hospital Universitário Clementino Fraga Filho, Universidade Federal do Rio de Janeiro, Rio de Janeiro, Brazil; Laboratório de Endocrinologia Experimental-LEEx, Instituto de Ciências Biomédicas, Universidade Federal do Rio de Janeiro, Rio de Janeiro, Brazil; Laboratório de Ultraestrutura e Biologia Tecidual, Instituto de Biologia Roberto Alcantara Gomes, UERJ, RJ; Laboratório Interdisciplinar de Pesquisas Médicas, IOC, Fiocruz, RJ, Brasil

**Keywords:** Chikungunya virus, Arbovirus, hyperglycemia, pathology

## Abstract

Chikungunya fever (CHIKF) is a re-emerging viral disease characterized by acute systemic manifestations and debilitating musculoskeletal symptoms that can persist after viral clearance. Although typically self-limiting in healthy individuals, clinical outcomes are significantly worsened in patients with pre-existing comorbidities, particularly diabetes mellitus (DM). Epidemiological data links DM to heightened CHIKF severity and a greater risk of developing chronic arthropathy, yet the mechanism underpinning this association remains poorly understood. In this study, we established an *in vivo* streptozotocin (STZ)-induced diabetic C57BL/6 mice as a model to investigate the impact of DM on CHIKV pathogenesis. STZ induces selective pancreatic β-cell destruction and persistent hyperglycemia. Diabetic animals infected with CHIKV exhibited aggravated joint inflammation, increased nociceptive sensitivity, and elevated serum markers of muscle and hepatic injury, including creatine kinase (CK) and alanine aminotransferase (ALT). Histopathological analyses revealed that CHIKV infection alone disrupted joint architecture. However, in the diabetic context, these alterations were significantly exacerbated, with enhanced inflammatory infiltrates, chondrocyte loss, osteocyte necrosis, and fibrotic remodeling. These results demonstrate that the diabetic metabolic environment profoundly amplifies CHIKV-induced tissue damage and impairs resolution of inflammation, offering a plausible mechanistic explanation for the poorest CHIKF outcomes observed in diabetic patients. Thus, this model provides a valuable platform for exploring the molecular drivers of CHIKF severity and chronicity, especially among DM patients, as well as for development of pharmacological tools to mitigate CHIKV-associated complications in metabolically vulnerable populations.

**Importance:** Chikungunya virus is responsible for a re-emerging disease that causes intense joint pain and long-lasting inflammation, especially in vulnerable individuals. People with diabetes are known to suffer more severe and persistent symptoms, but the biological reasons behind this have remained unclear. In this study, we used a diabetic mouse model to investigate how a high-glucose environment influences the course of Chikungunya virus infection. We found that diabetic mice experienced more intense joint damage, increased pain sensitivity, and signs of broader organ injury compared to non-diabetic animals. Microscopic analyses of tissues showed greater inflammation and structural damage in the joints of diabetic animals. These findings suggest that diabetes directly worsens the effects of Chikungunya virus infection by amplifying inflammation and delaying healing. This model helps explain why diabetic patients have worse outcomes and may assist in developing new treatments to protect high-risk populations from long-term complications of this infection.

## Introduction

Chikungunya fever (CHIKF) is an acute and debilitating viral illness caused by Chikungunya virus (CHIKV), a mosquito-borne *alphavirus* belonging to the *Togaviridae* family^1^. Since its initial isolation during outbreaks in the 1950s, CHIKV has emerged as a global health concern due to its capacity for rapid dissemination and recurrent urban epidemics^2,3^. CHIKV demonstrates a broad tissue tropism, infecting diverse cell types including keratinocytes, fibroblasts, myocytes, hepatocytes, chondrocytes, immune cells, among others^1^. As a result, the clinical manifestations of CHIKF are systemic and multifaceted, ranging from cutaneous and hepatic involvement to pronounced musculoskeletal and neurological symptoms^1,4^.

CHIKV infection is predominantly symptomatic, with only 15% of cases remaining subclinical^5^. The disease typically begins two to four days post-infection and manifests with an abrupt onset of high fever, maculopapular rash, conjunctivitis, retro-orbital pain, headache, nausea, and most notably, intense myalgia and arthralgia^6^. CHIKV also triggers a robust type I interferon response, which is critical for antiviral control but also contributes to lymphopenia and immunopathology^7,8^. Even after viral clearance, immune-mediated inflammation may persist, particularly in joint tissues, contributing to chronic arthritic symptoms in a significant proportion of patients.

Importantly, host comorbidities play a critical role in modulating CHIKF severity and progression^9^. Individuals with pre-existing metabolic or inflammatory conditions such as diabetes mellitus (DM), hypertension, obesity, or asthma exhibit increased susceptibility to atypical and more severe clinical outcomes^9^. Among these, DM is particularly notable, being associated with an approximately 20% increased risk of developing chronic post-CHIKF arthropathy^9–11^. The chronic nature of these complications severely impairs mobility and quality of life and represents a growing public health concern, especially in regions with high rates of metabolic disorders and CHIKV circulation.

DM is a heterogeneous metabolic disease defined by persistent hyperglycemia, arising from defects in insulin metabolism. Type 1 diabetes (T1DM) results from autoimmune destruction of pancreatic β-cells, while type 2 diabetes (T2DM) is characterized by insulin resistance and progressive β-cell dysfunction^12,13^. In both cases, impaired glucose regulation leads to widespread metabolic disturbances affecting multiple organ systems. Of particular relevance to CHIKF, insulin and its co-secreted peptide amylin are key regulators of musculoskeletal physiology. Insulin promotes bone remodeling by stimulating osteoblast activity and osteocalcin carboxylation^14,15^. Amylin contributes to joint and bone homeostasis by inhibiting osteoclast-mediated resorption, stimulating osteoblast and chondrocyte activity^16,17^. The deficiency or resistance to these hormones in DM disrupts bone integrity and joint homeostasis, leading to reduced trabecular density, structural fragility, and increased vulnerability to inflammatory damage^14,16^. Thus, the convergence of CHIKV infection with the altered metabolic and inflammatory milieu of DM may synergistically worsen tissue damage, prolong inflammation, and impair mechanisms of damage resolution, ultimately contributing to persistent joint pathology and more severe disease outcomes.

Given this clinical and mechanistic intersection, the present study aimed to establish and characterize an *in vivo* model of CHIKV infection under a pre-existing metabolic dysfunction. Male C57BL/6 mice were subjected to streptozotocin (STZ) treatment, a selective β-cell cytotoxic compound, to induce T1DM, which was confirmed by blood glucose monitoring. Wild-type (Control) and diabetic mice (STZ) were infected with CHIKV or mock-infected for 7 days long and, at the end, biological samples were collected for biochemical and histopathological analysis. Our data shows that diabetic infected mice exhibited increased levels of tissue damage, as creatine kinase (CK), lactate dehydrogenase (LDH), and alanine aminotransferase (ALT), when compared to other groups, suggesting a heightened muscle and liver injury. Histological evaluation further revealed that CHIKV infection alone induced inflammation, edema, and disorganization in joint tissues. However, in diabetic infected animals, these pathological features were significantly intensified, with marked increase in inflammatory infiltrates, tissue necrosis, and extracellular matrix disruption. Collectively, these findings provide compelling evidence that the interplay between viral infection and diabetic metabolic dysregulation amplifies tissue injury and impairs inflammatory resolution, offering a mechanistic explanation for the heightened susceptibility to chronic and severe outcomes of CHIKF observed in diabetic patients.

## Methodology

### Viral stocks and titration

Stocks of CHIKV (RJ-IB5 genotype ECSA, kindly gifted by professors Clarissa Damaso and Luciana Costa, Universidade Federal do Rio de Janeiro)^18^ were propagated by infecting *Aedes albopictus* mosquito cells (C6/36) cultured in L-15 medium supplemented with 5% fetal bovine serum (FBS) at 28°C in a B.O.D. incubator, as previously described^19^. Virus titration was performed in Vero-E6 following previously characterized protocols^20^.

### Animal model

This study was approved by the Ethics Committee on Animal Use in Research of the Federal University of Rio de Janeiro – UFRJ (protocols No. 041/21 and 115/21). Male C57BL/6 mice aged four to six weeks were randomly divided by body weight into 3 main groups: control, STZ 3 days, STZ 5 days. Mice were assessed for weight gain, casual blood glucose, and mechanical allodynia response. Animals were housed in a room with controlled temperature and a 12-hour light-dark cycle. Water and food were available *ad libitum*. Animals underwent an intraperitoneal administration protocol of multiple low doses of 50 mg/kg of STZ in 100 mM citrate buffer, pH 4.5, for 3 or 5 days. The control group received the drug vehicle (100 mM citrate buffer). During this period, mice were supplemented with 10% sucrose in water to prevent STZ-induced hypoglycemia. Ten days after the start of the protocol, blood glucose levels were monitored, and mice were considered diabetic when blood glucose exceeded 250 mg/dL for 3 consecutive readings^21^. Following DM confirmation, the groups were divided into subgroups to be inoculated with CHIKV or mock. Infected groups were inoculated with 10^6^ plaque-forming units (pfu) of CHIKV in the right hind paw and mock groups inoculated with the same volume of infection control. Seven days after infection, weight gain, casual blood glucose, and mechanical threshold were reassessed. Following these assessments, animals were anesthetized with a ketamine-xylazine cocktail (100 mg/kg ketamine and 10 mg/kg xylazine, intraperitoneally) to ensure deep anesthesia. Once full anesthesia was confirmed, blood was collected in heparin tubes via cardiac puncture for subsequent serum separation and biochemical analyses. Immediately thereafter, animals were cervically dislocated, and hindlimbs were extracted and fixated in 4% paraformaldehyde (PFA). Samples were further processed for histological and immunohistochemistry analysis as described below.

### Von Frey Test

Animals were acclimated for 1 hour and then subjected to sequential mechanical stimuli ranging from 0.008 g to 2.0 g using von Frey filaments. The stimulus was applied to both hind paws, and each filament was presented 5 times per paw with a 60-second interval between presentations. The mechanical threshold was defined as the filament that elicited the withdrawal response most frequently. Animals with a withdrawal threshold exceeding 2.0 g were excluded from the experimental groups (outliers). Mechanical sensitivity was measured on the day of infection and 7 days thereafter.

### Biochemical Analyses

Blood samples were obtained by cardiac puncture of mice using heparinized syringes (calcium-balanced lithium heparin). Next, samples were homogenized 5 times by inversion, transferred to microcentrifuge tubes and centrifuged at 2000 ×g for 10 min at room temperature. Plasma samples were transferred to new microcentrifuge tubes and stored at −20 °C until analysis.

Biochemical measurements such as creatinine kinase activity (CK) (Beckman OSR6179), lactate dehydrogenase activity (LDH) (Beckman OSR6127), alanine aminotransferase activity (ALT) (Beckman OSR6007) and triglycerides levels (Trig) (Beckman OSR60118) were quantitatively determined using spectrophotometric assays. The very low-density lipoprotein cholesterol levels (VLDL-C) were estimated from Trig levels using the following formulae: VLDL-C = Trig / 5.

Assays were performed in a clinical analyzer (Beckman AU5800 and AU680 autoanalyzers, Beckman Coulter Diagnostics, Brea, CA) at HUCFF-UFRJ.

### Histology and immunohistochemistry analysis

Following fixation, hindlimbs were subjected to decalcification in a 20% ethylenediaminetetraacetic acid (EDTA) solution, adjusted to pH 7.4, and maintained at 4°C under constant gentle agitation. The decalcification process was carried out for two weeks, with the EDTA solution being refreshed every two to three days to ensure optimal chelation efficiency and complete removal of mineral content. Samples were then rinsed in PBS to remove residual EDTA and dehydrated through a graded ethanol series (70%, 80%, 90%, 95%, and 100%), followed by clearing in xylene. Subsequently, tissues were embedded in paraffin blocks and sectioned with a microtome into 4µm sections. To evaluate general tissue architecture, inflammatory infiltrates, and structural alterations in joint compartment, sections were stained with hematoxylin and eosin (H&E). For specific detection of cellular or molecular markers, antigen retrieval was performed using citrate buffer at pH 6.0 with incubation period of 20 min at 60°C. Endogenous peroxidase activity was blocked using 0.5% hydrogen peroxide (H_2_O_2_) and non-specific binding of the polyclonal antibodies was blocked by incubation 5% BSA. After, sections were incubated with antibodies, and these reactions were amplified using a biotin–streptavidin system (Vector Labs, USA). We used anti-E-CHIKV, anti-CD8, anti-TNF-α, anti-IL-1b, (dilution 1:100, Santa Cruz Biotechnology, EUA) antibodies. Immunoreactive products were visualized using diaminobenzidine (DAB) reagent (Vector) and counter stained with hematoxylin. Images were analyzed under a light microscope equipped with a CCD camera (Olympus BX53 with camera Olympus DP72, Nagano, Chubu, Japan) using the software Image-Pro Plus 7.0 program (Media Cybernetics, Silver Springs, Maryland, EUA).

### Morphometry

The slices were stained with safranine O-fast green for cartilage analysis and then observed under a light microscope equipped with a CCD camera (Olympus BX53 with camera Olympus DP72, Nagano, Chubu, Japan). Five random fields were obtained from each slide of animal-group. The Image-Pro Plus 7.0 program (Media Cybernetics, Silver Springs, Maryland, EUA) was used to quantify the cartilage area with a magnification of 20x.

### Statistical analysis

Data were analyzed with GraphPad prism software v 6.0 (San Diego, California, USA) using analysis of one-way ANOVA. The differences between groups were considered statistically significant when values of p <0.05. a-different from the control and b-different from the STZ.

## Results

### CHIKV infection promoted heightened paw swelling in diabetic mice but did not influence host weight and glycemia

Several well-established animal models are available for the study of diabetes, with the STZ and high-fat diet (HFD)-induced models being among the most frequently employed^22^. The STZ model is widely used to replicate T1DM, as this drug selectively destroys pancreatic β-cells, resulting in rapid and sustained hyperglycemia due to insulin deficiency^23^. On the other hand, the HFD model is primarily used to mimic T2DM, where prolonged consumption of a lipid-rich diet promotes obesity, insulin resistance, and progressive β-cell dysfunction^24^. Both models are valuable tools in diabetes research, however, they differ in their timelines and pathophysiological characteristics. The STZ model allows for a faster, more uniform onset of diabetic symptoms, making it suitable for studies requiring rapid disease induction. Conversely, the HFD model more accurately recapitulates the complex, gradual development of metabolic syndrome, although it demands a longer experimental timeline. Given these considerations, we selected the STZ-induced model for our study to efficiently investigate the impact of CHIKV infection in the context of DM. Thus, male C57BL/6 mice aged four to six weeks were divided in three main groups: control, STZ 3 days or STZ 5 days. Mice were treated with multiple low doses of STZ for 3 or 5 days and were considered diabetic when glycemia exceeded 250 mg/dL in 3 consecutive readings. Mice that received the 3-day STZ treatment protocol did not develop hyperglycemia and were therefore excluded from subsequent analyses (data not shown). In contrast, animals treated with STZ for 5 consecutive days exhibited sustained hyperglycemia, with blood glucose levels exceeding 300 mg/dL, and were classified as diabetic. Following the successful induction of diabetes, experimental groups were randomly subdivided into two subgroups: mock-infected or CHIKV-infected. For infection, viral and mock inoculum were freshly prepared and administered subcutaneously into the right hind paw (plantar region). Animals were then monitored daily over the course of one week to capture clinical signs of disease. Infected groups exhibited paw swelling, with the STZ-treated mice displaying exacerbated edema compared to non-diabetic controls (Fig. 1A). However, no significant changes in body weight or blood glucose levels were observed in either group following CHIKV infection (Fig. 1B and 1C). These findings indicate that while CHIKV infection provoked a localized inflammatory response, characterized by local edema, particularly intensified under diabetic conditions, it did not induce glycemia and weight alterations within the monitored timeframe.

**Figure 1.**
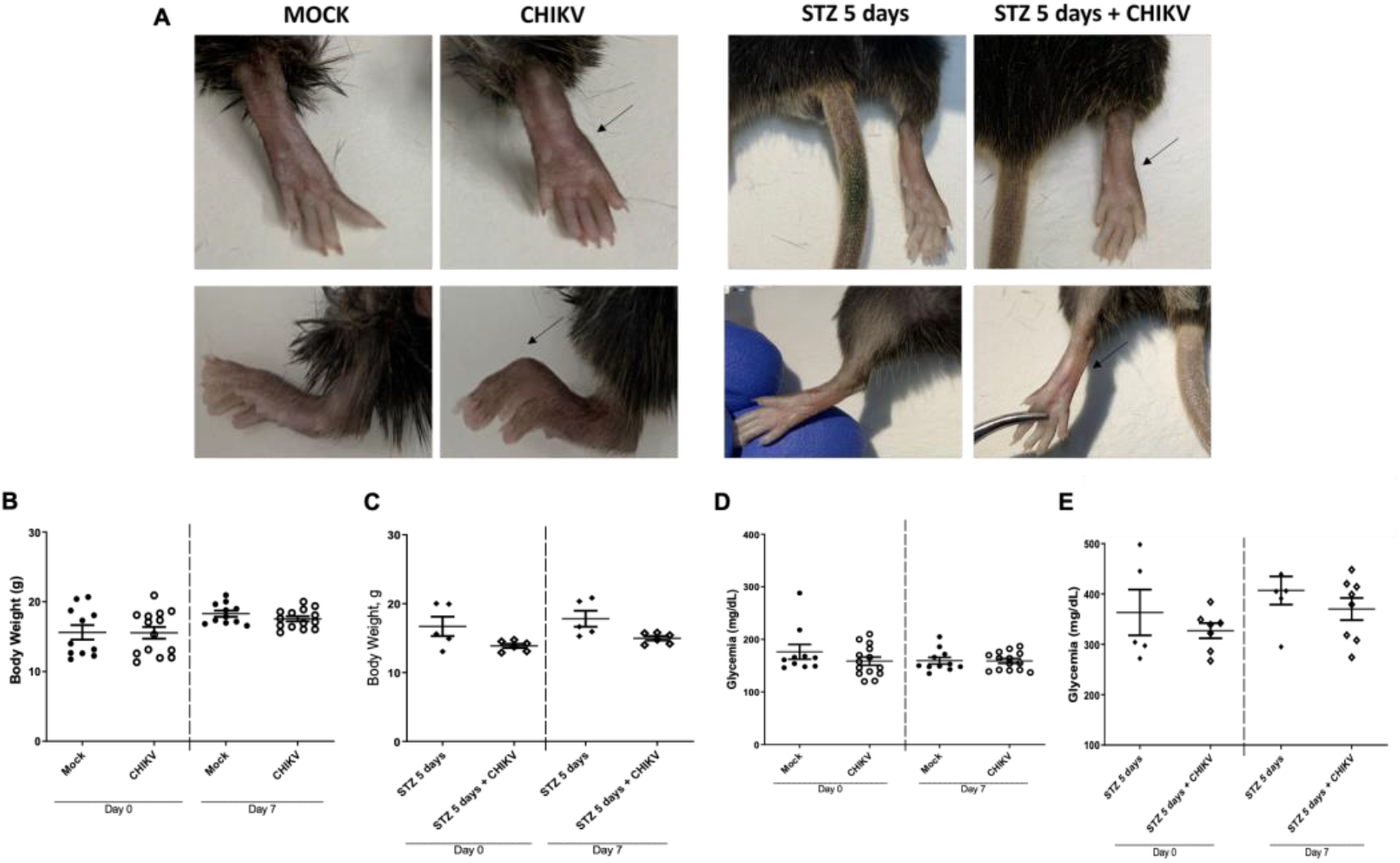
Assessment of clinical parameters following CHIKV infection in diabetic and non-diabetic mice. C57BL/6 mice with streptozotocin (STZ)-induced diabetes and controls were infected with CHIKV and monitored daily over 7 days. (A) Paw edema was observed to evaluate localized inflammation (arrows). (B-C) Body weight was recorded periodically to assess systemic health status. (D-E) Blood glucose concentrations were measured using glucometry to monitor glycemic changes post-infection. Data are presented as mean ± SEM.

### CHIKV infection induced alterations in muscle and hepatic tissues markers

To further investigate whether CHIKV infection contributes to systemic metabolic disturbances, we analyzed a panel of serum biochemical markers reflective of tissue integrity and metabolic status. CK and ALT, enzymes that indicate muscle and liver damage, respectively^25^, were significantly elevated only in the CHIKV-infected STZ-treated group (Fig. 2A and B), suggesting synergistic tissue stress when viral infection co-occurs with diabetes. In contrast, LDH, a broader marker of cellular injury, was elevated in both CHIKV-infected groups regardless of diabetic status (Fig. 2C), indicating a general response to viral infection. On the other hand, serum levels of triglycerides and VLDL, often altered during metabolic dysregulation, remained unchanged across all groups following CHIKV infection (Fig. 2D and E). Collectively, these data suggest that while CHIKV infection alone can induce mild systemic cellular stress, the presence of diabetes amplifies tissue-specific damage, particularly in muscle and liver.

**Figure 2.**
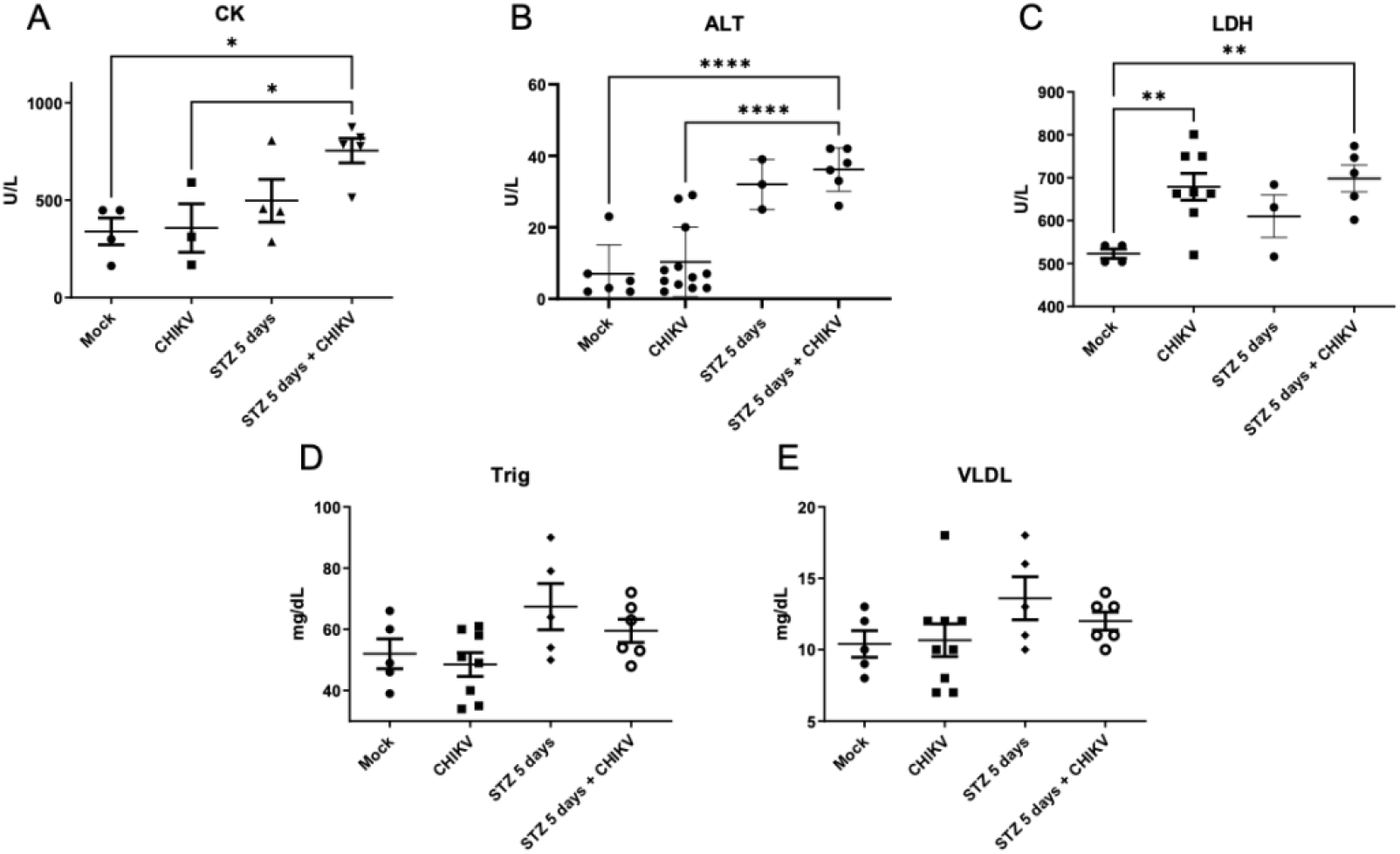
Serum biochemical analyses in CHIKV-infected diabetic and non-diabetic mice. Enzymatic activity of (A) creatine kinase (CK) and (B) alanine aminotransferase (ALT) were measured to evaluate tissue damage. (C) Lactate dehydrogenase (LDH) activity was quantified as a marker of general cellular injury. Serum (D) triglycerides and (E) very-low-density lipoprotein cholesterol (VLDL-C) concentrations (mg/dL) were assessed to determine systemic metabolic alterations. Data are presented as mean ± SEM (n=5).

### CHIKV infection interferes with pain perception in DM mice

Paw swelling, resulting from localized inflammatory responses, can markedly affect the interpretation of pain signals^26^. The accumulation of inflammatory exudate and immune cell infiltration leads to tissue edema, which increases local pressure and disrupts the mechanical integrity of the skin, connective tissue, and underlying nociceptive structures^27^. This swelling can have dual effects on pain perception: on one hand, it may sensitize peripheral nociceptors, lowering their activation threshold and enhancing pain responses to normally innocuous stimuli; on the other hand, excessive edema may mechanically buffer or disperse pressure stimuli^27,28^. To evaluate whether CHIKV-induced paw swelling interferes with pain sensitivity, we assessed mechanical allodynia using Von Frey filaments across our experimental groups. Mice in the infected control group showed no significant changes in mechanical sensitivity immediately following CHIKV inoculation or 7 days post-infection (Fig. 3A and B), suggesting that infection alone did not alter nociceptive thresholds in non-diabetic animals. However, in the STZ-treated group, a marked increase in sensitivity to Von Frey stimuli were observed 7 days post-infection (Fig. 3C and D). These animals exhibited enhanced responsiveness to lower filament forces, indicative of heightened pain perception (Fig. 3D). This indicates that the combination of diabetes and CHIKV infection synergistically amplifies peripheral nociceptive signaling, potentially due to a pro-inflammatory environment that exacerbates neuroinflammatory responses and sensitizes pain pathways.

**Figure 3.**
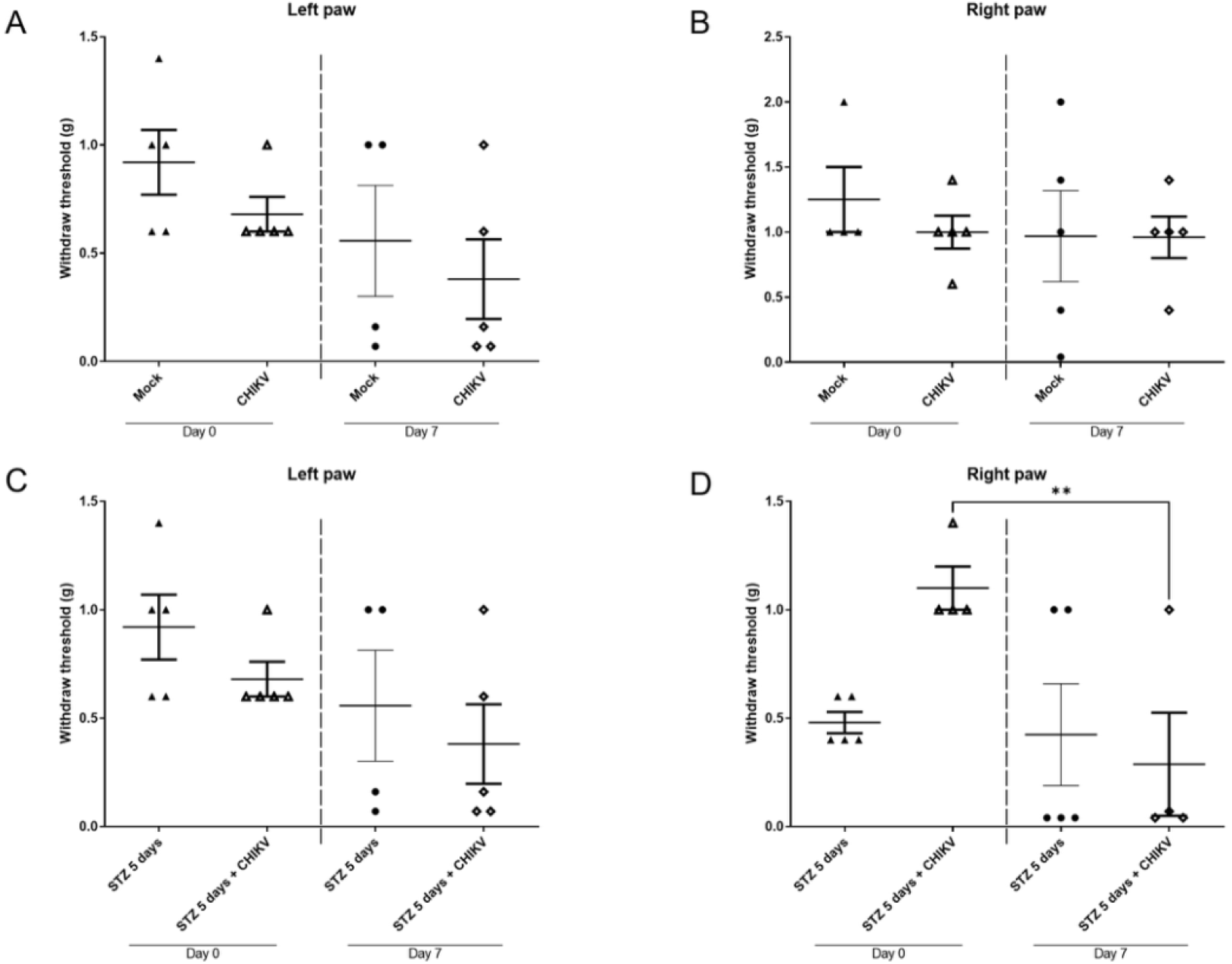
Evaluation of mechanical nociception following CHIKV infection in diabetic and non-diabetic mice. Mechanical allodynia was assessed using Von Frey filaments. (A, B) Nociceptive thresholds were measured in control-infected mice immediately after infection and 7 days post-infection. (C, D) Nociceptive sensitivity was similarly assessed in STZ-treated diabetic mice at the same time points. Data are presented as mean ± SEM.

### CHIKV-infected diabetic mice presented exacerbated tissue damage and loss of structural integrity and architectural organization

To gain a more in-depth understanding of how diabetes influences CHIKV-induced joint pathology, we performed comprehensive histological evaluations of the hindlimb joints. In the Mock group, all examined structures — including the articular cartilage, growth plate, metaphysis, subchondral bone, and meniscus — displayed preserved architecture, with well-organized tissue layers and no signs of degeneration or inflammation (Fig. 4A-B). In contrast, CHIKV-infected mice exhibited pronounced histopathological alterations. Notable features included thinning of the articular cartilage, irregular joint space partially infiltrated by marrow-like tissue, and disorganized trabecular bone. These animals also showed reduced bone matrix deposition, necrotic osteocytes, fibrotic areas indicative of aberrant tissue remodeling, and clear chondrocyte loss within cartilage layers — hallmarks of osteoarthritic-like degeneration (Fig. 4C-D). Interestingly, STZ treatment alone was sufficient to provoke structural disturbances in joint architecture, such as partial matrix disorganization and early signs of cartilage erosion. However, these changes were markedly exacerbated in animals subjected to both STZ treatment and CHIKV infection (Fig. 4E-H). In the CHIKV-STZ group, we observed a profound loss of joint matrix and a significant reduction in viable chondrocytes, suggesting that the combination of pancreatic β-cell damage and viral infection synergistically intensifies joint destruction (Fig. 4G-H). Furthermore, immunohistochemistry analysis revealed a robust infection and inflammatory profile in infected joints (Fig. 5A-H). CD8+ T lymphocyte infiltration (Fig. 6A-H), along with elevated staining for pro-inflammatory cytokines TNF-α and IL-1β (Fig. 7A-H), was observed in all infected groups but was absent in the mock controls. Notably, both CHIKV-infected groups exhibited significant reductions in the cartilaginous area of the articular surface and epiphyseal growth plate, pointing to impaired cartilage homeostasis and accelerated degradation (Fig. 8A-E). Collectively, these findings demonstrate that CHIKV infection alone can initiate joint degeneration. However, when combined with the diabetic milieu, the pathological outcomes are markedly intensified. The observed lesions closely resemble osteoarthritic damage, reinforcing the concept that metabolic comorbidities such as DM potentiate CHIKV-induced musculoskeletal complications.

**Figure 4.**
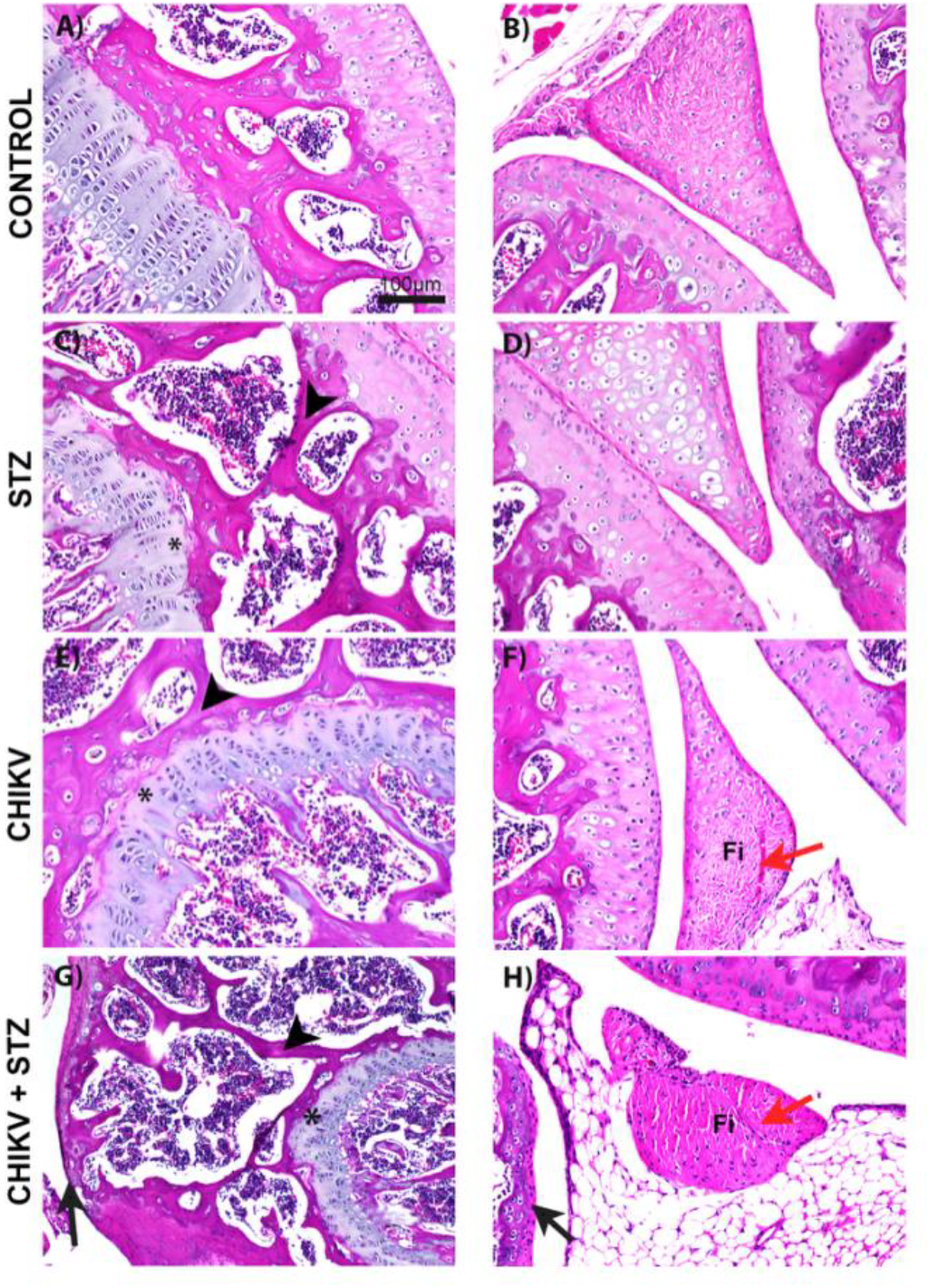
Histopathological assessment of hindlimb joints in diabetic and non-diabetic mice following CHIKV infection. Representative micrographs illustrate joint architecture, including articular cartilage, growth plate, metaphysis, subchondral bone, and meniscus. (A, H) Mock controls showing preserved joint structures, while CHIKV-infected groups exhibiting alterations in cartilage thickness, joint space, trabecular organization, bone matrix deposition, and chondrocyte viability (asterisks). Decrease and necrosis of chondrocytes in the cartilage in the area of endochondral ossification, in the zone of columnar and hypertrophic cartilage (arrowhead). Areas of probable fibrosis (Fi). Chondrocytes undergoing cell death process (Red arrows). Loss of articular matrix and chondrocytes (Black arrows).

**Figure 5.**
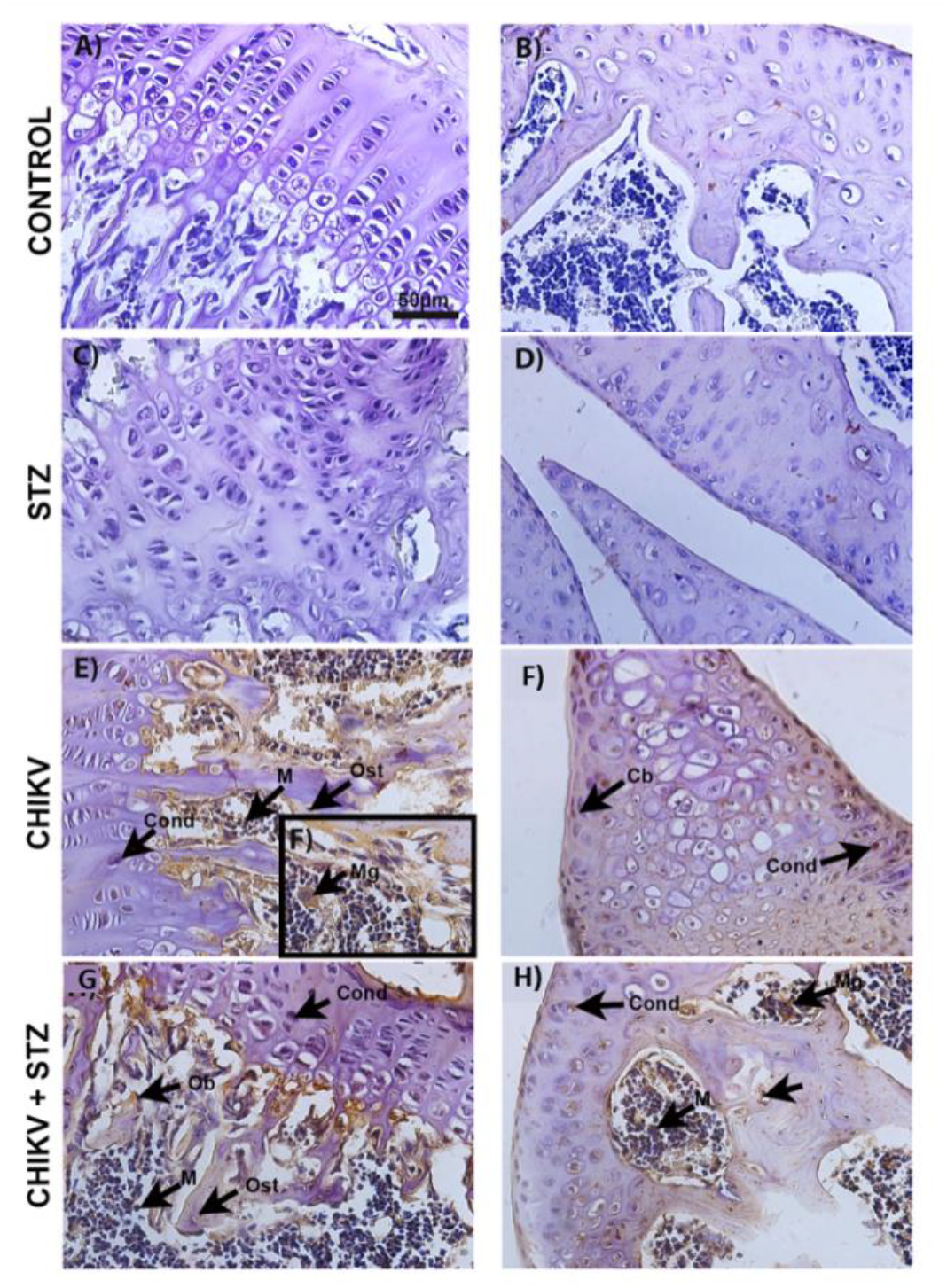
Confirmation of virus presence in infected groups, CHIKV and CHIKVSTZ. Virus presence was evidenced by specific labeling in various cell types: chondroblasts (Cb), chondrocytes (Cond), bone marrow cells (M), megakaryocytes (Mg), osteoblasts (Ob), and osteocytes (Ost).

**Figure 6.**
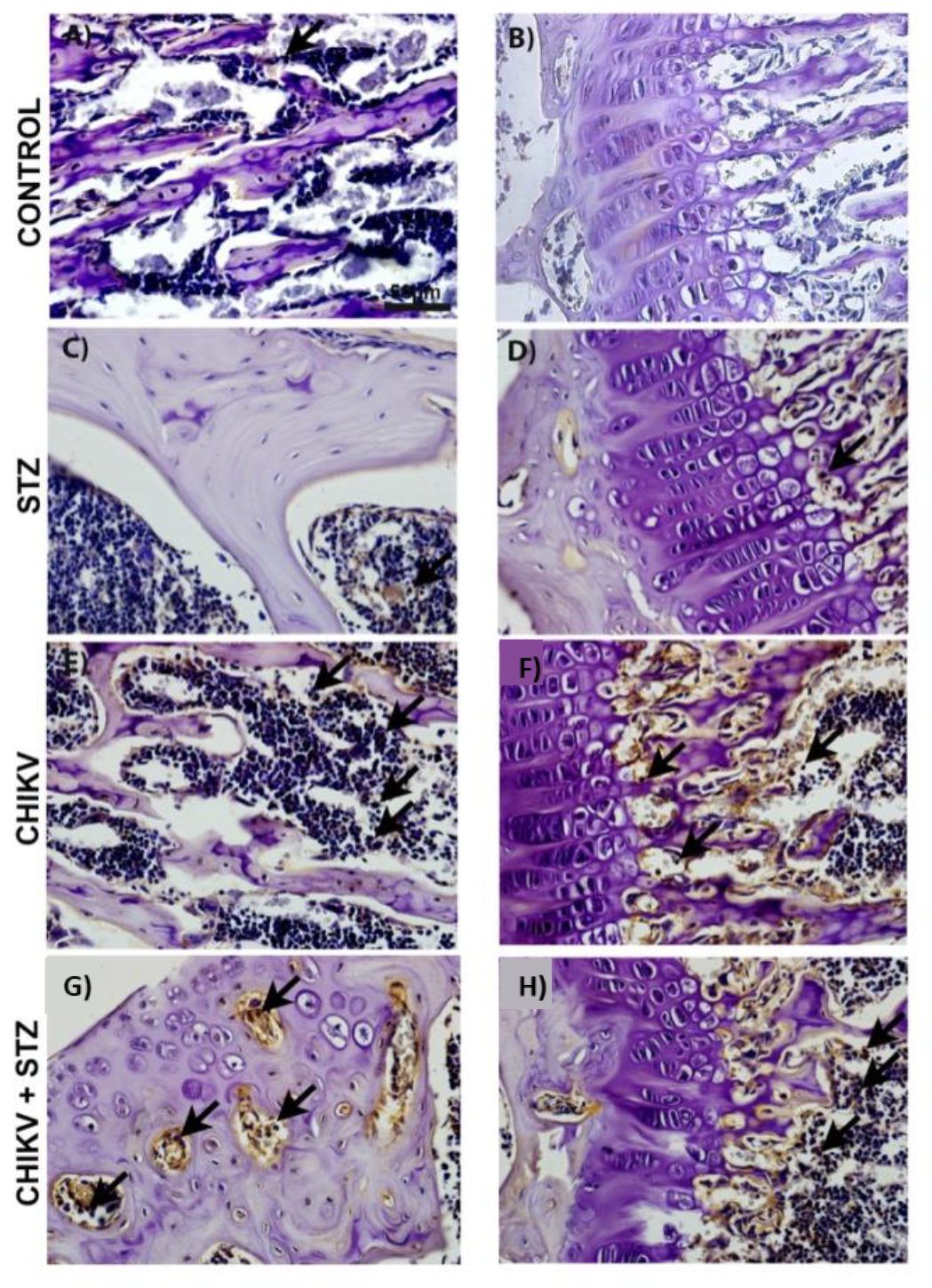
CD8+ T cell infiltration confirms immune activation in infected tissues. Immunohistochemical staining for CD8+ T lymphocytes reveals increased inflammatory cell infiltration. Arrows indicate CD8+ positive cells within affected tissue regions.

**Figure 7.**
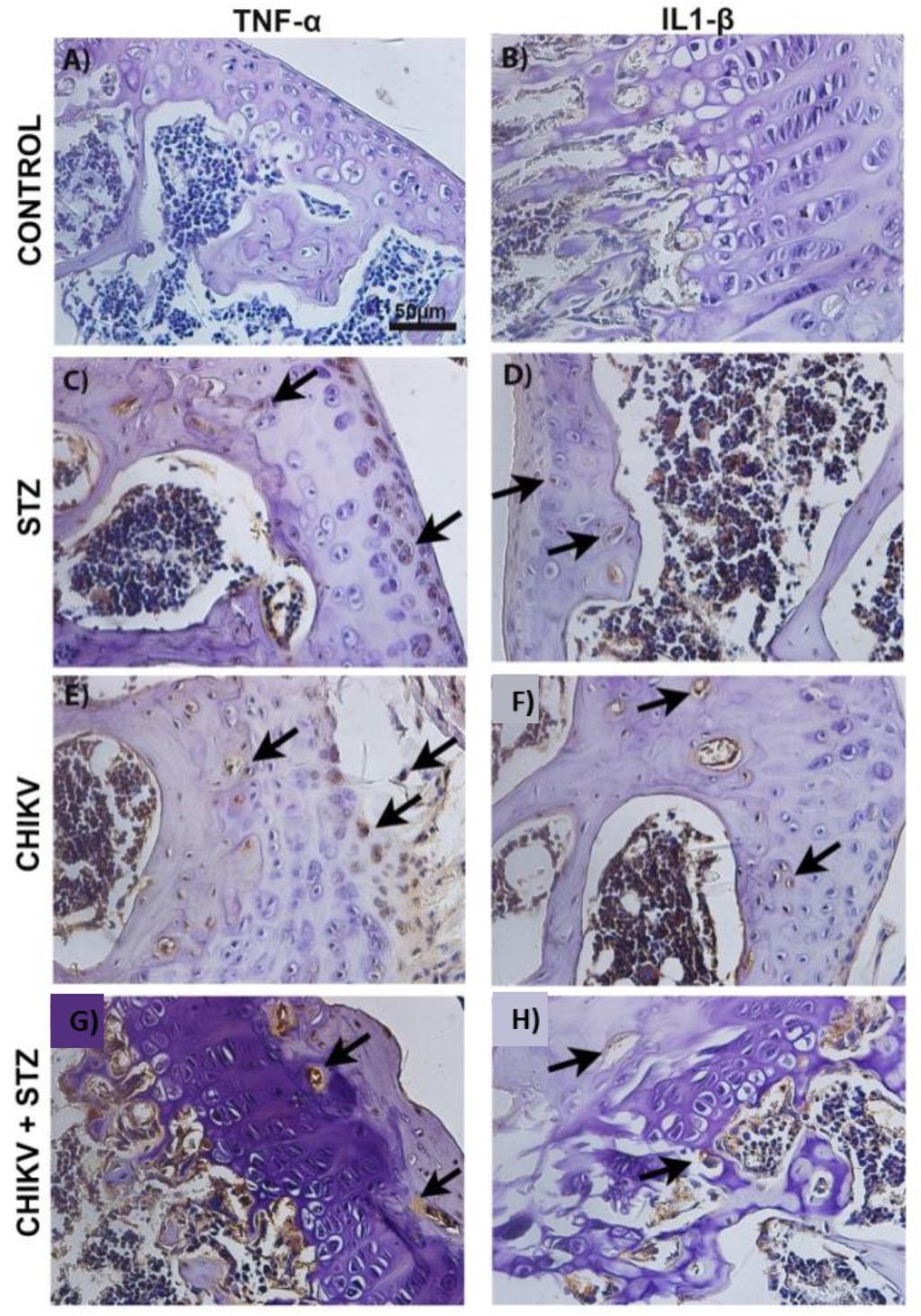
Increased expression of TNF-α and IL-1β confirms tissue inflammation. Immunohistochemical staining for TNF-α and IL-1β (A–H) reveals elevated cytokine levels in affected tissues. Arrows indicate positive staining sites, confirming the presence of inflammatory mediators.

**Figure 8.**
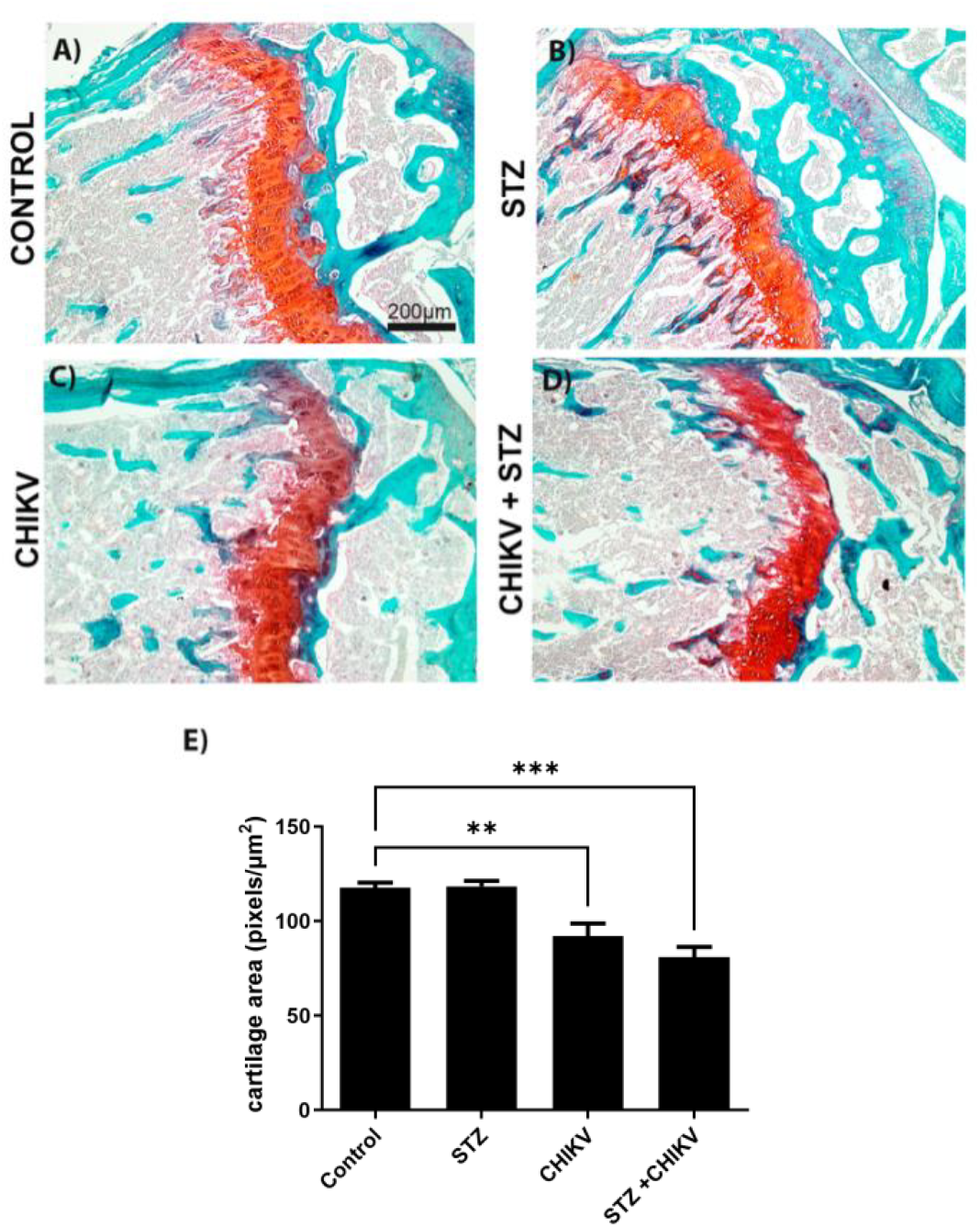
Reduced cartilaginous area and decreased chondrocyte numbers indicate cartilage degeneration in infected groups. Quantitative analyses of the cartilaginous area and chondrocyte counts were performed to assess cartilage integrity and degeneration. Special staining with Safranin for cartilage and quantification of cartilage area by morphometry. It is possible to observe a decrease in red-stained area, which in special staining highlights the cartilaginous area, both in the joint and epiphyseal disk. Data are presented as mean ± SEM.

## Discussion

CHIKF is a debilitating mosquito-borne disease characterized by the abrupt onset of high fever, skin rash, and incapacitating polyarthralgia, which can persist for months or even years after the initial infection^1^. Although typically self-limiting and non-fatal in immunocompetent individuals, an increasing number of reports have documented atypical and severe cases, especially in patients with pre-existing comorbidities such as DM, hypertension, and cardiovascular diseases^9^. These pre-existing chronic conditions have been associated with heightened risk of hospitalization, prolonged symptomatology, and chronic arthralgia^9^. Despite growing clinical awareness of these associations, the mechanisms through which metabolic dysfunctions such as DM exacerbate the clinical outcomes of CHIKF remain insufficiently understood.

Given the rising global burden of both arboviral infections and metabolic syndromes, we focused our study on elucidating how diabetes modulates the pathogenesis of CHIKV infection. Therefore, we employed a STZ-induced mouse model of DM. STZ is a β-cell-specific cytotoxin that induces sustained hyperglycemia by destroying insulin-producing cells in the pancreas, thus mimicking key features of T1DM^23^. Our data demonstrates that pre-existing DM significantly worsens the pathological effects of CHIKV infection, especially in terms of joint integrity and pain perception. In line with previous studies using non-diabetic mice^29^, we observed that CHIKV infection alone induces localized paw swelling, a hallmark of peripheral inflammation. Importantly, however, no substantial systemic metabolic disturbances — such as alterations in blood glucose or body weight — were detected during the early stages of infection in these animals. This observation supports the notion that, in otherwise healthy hosts, CHIKV infection tends to remain localized and does not necessarily perturb systemic homeostasis. In contrast, diabetic mice showed marked exacerbation of CHIKV-induced joint damage and pain. Despite the infection not further altering glycemia or weight in STZ-treated animals, histopathological and behavioral assessments revealed extensive joint destruction, increased immune infiltration, and intensified mechanical allodynia. These findings are consistent with reports from clinical cohorts, which demonstrated that diabetic patients with CHIKF experience more intense and prolonged joint symptoms compared to non-diabetic individuals^9,10^.

Von Frey testing in our model revealed that CHIKV-infected diabetic animals exhibited greater sensitivity to mechanical stimuli, even in the presence of similar paw swelling as their non-diabetic counterparts. This suggests that hyperglycemia and the chronic inflammatory milieu characteristic of diabetes may prime the peripheral nervous system for heightened pain responses. Supporting this, our immunohistochemistry analyses showed increased tissue expression of pro-inflammatory cytokines (TNF-α and IL-1β) and CD8+ T cell infiltration, both of which are implicated in neuroinflammation and nociceptive sensitization^30^. These results provide mechanistic insight into how metabolic inflammation may synergize with viral-induced damage to worsen clinical outcomes.

Histological examination of joint architecture strongly reinforced the notion of synergistic tissue damage resulting from the interplay between CHIKV infection and DM.

In non-diabetic animals, CHIKV infection alone was sufficient to induce significant alterations in joint structure, including thinning of the articular cartilage, reduction of bone matrix density, and localized fibrotic remodeling. These features closely resemble early-stage osteoarthritic lesions^31,32^ and are consistent with previous descriptions of CHIKV-induced arthritis in both animal models and human cases^33,34^. However, these pathological changes were markedly more pronounced in diabetic animals, suggesting that pre-existing metabolic dysfunction amplifies viral-induced joint degeneration.

Notably, even mock-infected STZ-treated mice exhibited signs of joint compromise, including mild cartilage erosion, subchondral bone irregularities, and signs of extracellular matrix degradation. These observations are consistent with the well-documented deleterious effects of STZ on skeletal tissues^35–37^. STZ-induced diabetes leads to systemic metabolic disturbances, including hyperglycemia, insulin deficiency, and oxidative stress, all of which are known to impair bone remodeling, enhance osteoclastic activity, and suppress chondrocyte viability. Previous studies have shown that STZ exposure promotes osteopenia, increases bone resorption, and disrupts cartilage matrix homeostasis, mechanisms that are likely to contribute to the joint abnormalities observed in mock-STZ animals^35–37^. Importantly, when CHIKV infection occurred on this compromised metabolic background, the resulting joint pathology was dramatically exacerbated. Diabetic CHIKV-infected mice displayed widespread chondrocyte apoptosis, extensive osteocyte necrosis, pronounced trabecular disorganization, and severe fibrotic infiltration within the joint capsule and synovial tissue. These alterations point to a breakdown of both cartilage and bone homeostasis, likely driven by heightened inflammatory responses and impaired tissue repair capacity in the diabetic milieu. This compounded joint degeneration may reflect the convergence of two pathogenic processes: the direct inflammatory and cytopathic effects of CHIKV on musculoskeletal tissues, and the chronic metabolic stress imposed by diabetes, which diminishes the ability of these tissues to respond and adapt to injury. The observed reduction in articular cartilage thickness, the collapse of epiphyseal architecture, and the presence of osteolytic lesions in CHIKV-STZ animals strongly suggest an acceleration of joint degradation pathways reminiscent of advanced arthropathies.

Together, these findings suggest that pre-existing diabetes not only exacerbates CHIKV-induced joint inflammation and degeneration but also intensifies pain-related behaviors and subclinical tissue stress (Fig. 9). These results emphasize the clinical importance of considering metabolic comorbidities when evaluating CHIKV disease progression, especially in regions where both diabetes and arboviral infections are highly prevalent. While the STZ model used here has provided valuable insights into how insulin deficiency and hyperglycemia contribute to CHIKV pathogenesis, it does not fully capture the multifactorial nature of T2DM, which involves insulin resistance, chronic low-grade inflammation, and altered lipid metabolism. Therefore, to gain a more comprehensive understanding of how diverse forms of metabolic dysfunction influence viral outcomes, future studies should incorporate complementary models such as HFD-induced insulin resistance or genetically modified models of T2DM. These approaches will be crucial for discerning whether the mechanisms uncovered here are specific to β-cell loss or reflect a broader susceptibility of metabolically compromised tissues to arboviral injury.

**Figure 9.**
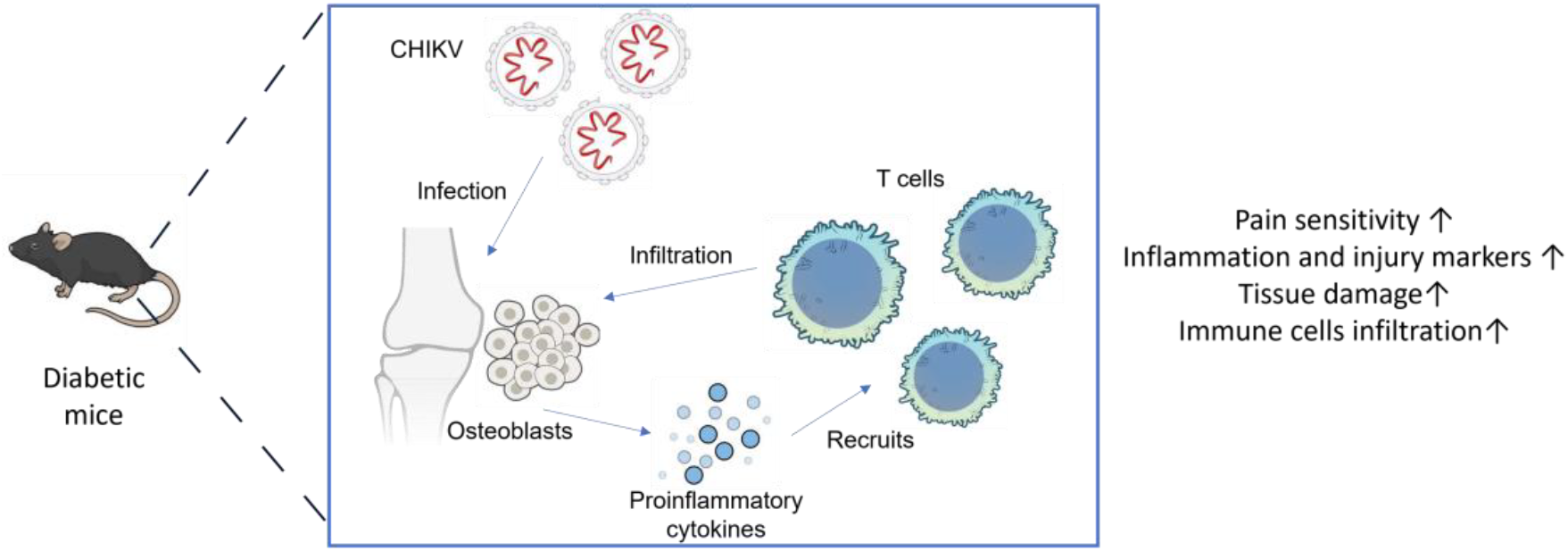
Schematic representation of the interplay between diabetes mellitus and Chikungunya virus (CHIKV) infection in exacerbating joint pathology. The model illustrates how streptozotocin (STZ)-induced hyperglycemia creates a pro-inflammatory metabolic environment that amplifies CHIKV-induced localized inflammation and tissue damage. In diabetic mice, CHIKV infection leads to exacerbated paw edema, heightened nociceptive sensitivity, increased muscle and liver injury markers, and severe joint histopathology characterized by cartilage degradation, bone remodeling, and immune cell infiltration. This synergy results in aggravated musculoskeletal complications, highlighting the importance of metabolic comorbidities in modulating viral disease outcomes.

